# Remapping the cognitive and neural profiles of children who struggle at school

**DOI:** 10.1101/295725

**Authors:** Duncan E. Astle, Joe Bathelt, the CALM team, Joni Holmes

## Abstract

Our understanding of learning difficulties largely comes from children with specific diagnoses or individuals selected from community/clinical samples according to strict inclusion criteria. Applying strict exclusionary criteria overemphasizes within-group homogeneity and between-group differences, and fails to capture comorbidity. Here we identify cognitive profiles in a large heterogeneous sample of struggling learners, using unsupervised machine learning in the form of an artificial neural network. Children were referred to the Centre for Attention Learning and Memory (CALM) by health and education professionals, irrespective of diagnosis or comorbidity, for problems in attention, memory, language, or poor school progress (n=530). Children completed a battery of cognitive and learning assessments, underwent a structural MRI scan, and their parents completed behaviour questionnaires. Within the network, we could identify four groups of children: i) children with broad cognitive difficulties, and severe reading, spelling and maths problems; ii) children with age-typical cognitive abilities and learning profiles; iii) children with working memory problems; and iv) children with phonological difficulties. Despite their contrasting cognitive profiles, the learning profiles for the latter two groups did not differ: both were around 1 SD below age-expected levels on all learning measures. Importantly a child’s cognitive profile was not predicted by diagnosis or referral reason. We also constructed whole-brain structural connectomes for children from these four groupings (n=184), alongside an additional group of typically developing children (n=36), and identified distinct patterns of brain organisation for each group. This study represents a novel move towards identifying data-driven neurocognitive dimensions underlying learning-related difficulties in a representative sample of poor learners.

**Author Note:** The Centre for Attention Learning and Memory (CALM) research clinic is based at and supported by funding from the MRC Cognition and Brain Sciences Unit, University of Cambridge. The Principal Investigators are Joni Holmes (Head of CALM), Susan Gathercole (Chair of CALM Management Committee), Duncan Astle, Tom Manly and Rogier Kievit. Data collection is assisted by a team of researchers and PhD students at the CBSU. This currently includes: Sarah Bishop, Annie Bryant, Sally Butterfield, Fanchea Daily, Laura Forde, Erin Hawkins, Sinead O’Brien, Cliodhna O’Leary, Joseph Rennie, and Mengya Zhang. The authors wish to thank the many professionals working in children’s services in the South-East and East of England for their support, and to the children and their families for giving up their time to visit the clinic.

**Research Highlights:**

- first study to apply machine learning to understand heterogeneity in struggling learners
- large sample of struggling learners that includes children with multiple difficulties
- rich phenotyping with detailed behavioural, cognitive, and neuroimaging assessments

---

Prevalence rates of developmental disorders linked with learning difficulties, including attention deficit hyperactivity disorder (ADHD), dyslexia, dyscalculia and specific language impairment (SLI), range from 3–8 % (American Psychiatric Association, 2013; Norbury et al., 2016; Polanczyk et al., 2014; Shalev and Gross-Tsur, 2001). But the number of children who struggle at school is far higher. In the UK for example, around 30% of the school population fail to meet expected targets in reading or maths at age 11 (DfE, 2017). The long-term outcomes for children who struggle at school include continued educational underachievement, poor mental health (Roeser and Eccles, 2000) and underemployment (de Beer et al., 2014; Bynner and Parsons 2005).

Our understanding of the causes of learning difficulties comes largely from studying children with a specific diagnosis (e.g. ADHD or SLI) or those selected from community or clinical samples on the basis of strict inclusion criteria (e.g. children with poor reading skills, but age-typical IQ and maths abilities). Most studies recruit children with “pure” problems (e.g. children with ADHD without comorbid dyslexia, or children with maths problems in the absence of reading problems or low IQ). There are practical advantages to this approach: it outlines clear criteria to inform practitioner decision-making about primary areas of weakness that can be used to identify intervention options.

However, this approach can fail to accommodate the high rates of comorbidity within developmental disorders (Kotov et al., 2017; Coghill & Sonuga-Barke, 2012) and learning-related difficulties (e.g., Angold et al., 1999). Over 80% of children with ADHD meet criteria for at least one additional diagnosis (e.g., Faraone et al., 1998; Willcutt & Pennington, 2000) and 15-45% have co-occurring reading difficulties (e.g., Biederman et al., 1991; Faraone et al., 1993; Semrud-Clikeman et al., 1992). Reading difficulties also co-occur 50% of the time with maths (Moll et al., 2014) or language problems (McArthur et al., 2000).

Using strict exclusionary criteria also overemphasizes similarities within groups, and the distinctiveness between groups (Kotov et al., 2017; Coghill & Sonuga-Barke, 2012). It is widely documented that symptoms vary between children with the same diagnosis. For example, performance on cognitive tasks within ADHD groups is notoriously variable (Castellanos et al., 2005; Nigg et al., 2005). Symptoms also co-occur across groups. For example, symptoms of inattention are common in children with poor literacy and maths skills (Hart et al., 2010; Loe & Feldman, 2007; Zentall, 2007), ADHD, autism spectrum disorder (ASD; Rommelse, Geurts, Franke, Buitelaar & Hartman, 2011), SLI (Duinmeijer, Jong & de Scheper, 2012), and dyslexia (Willcutt & Pennington, 2000; Germano, Gagliano, & Curatolo, 2010). Finally, this approach of selectively grouping children does not capture the majority of struggling learners – they often do not receive a diagnosis or are characterised by complex and comorbid difficulties that would rule them out of studies with strict inclusion criteria.

For these reasons a number of researchers have advocated empirically-based quantitative classification systems (Archibald et al., 2013; Coghill & Sonuga-Barke, 2012; Ramus et al., 2013; Sonuga-Barke & Coghill, 2014), although few studies have done this. The aim of this approach is to move away from identifying highly selective discrete groups and instead focus on identifying continuous dimensions that distinguish individuals and can be used as potential targets for intervention. Dimensions are derived through data-driven explorations of the data, with no a priori assumptions about group membership. For example, factor analysis, a statistical method that groups variables based on shared variance, is used most commonly to derive underlying dimensions from sets of symptoms or measures (e.g. Kotov et al., 2017). This technique has been used to identify dimensions of phonological and non-phonological skills in children with diagnosed SLI and dyslexia (Ramus et al., 2013) and separate latent constructs for inattention and hyperactivity in children with ADHD (Martel, von Eye & Nigg, 2010). An alternative approach, as yet rarely used, is to cluster children together according to shared profiles based on empirical data. In turn this can be used to inform classification systems, and consequently treatment approaches. Clustering algorithms have been used to identify groups of children with distinct learning (Archibald, Cardy, Joanisse & Ansari, 2013) and behavioural profiles (Bathelt, Holmes, the CALM Team & Astle, 2017).

In this study we use a different data-driven approach – machine learning. Machine learning methods have rarely been applied to understanding developmental disorders (e.g. Fair et al., 2012). Typical applications use supervised machine learning (Peng et al., 2013) in which the algorithm attempts to learn about pre-defined categories of children. Here we use an unsupervised learning approach whereby the algorithm attempts to learn about the structure of the data itself rather than which data correspond to pre-defined groups. Specifically, we used Self Organising Maps (SOMs; Kohonen, 1989), a type of artificial neural network. Due to their efficacy in visualising multidimensional data, SOMs have been successfully applied to a variety of tasks including textual information retrieval (Lin, 1991), the interpretation of gene expression data (Tamayo, 1999), and ecological community modelling (Giraudel, 2001). SOMs use an algorithm that projects the original data from a multidimensional input space onto a two-dimensional grid of nodes called a ‘map’, while preserving topographical information. This produces an inter-variable representational space, wherein the geometric distance between nodes corresponds to the degree of similarity in the input data. Within the current context, input data are individual children from our sample. The map will represent the cognitive profiles of the children; the closer the children are represented within the map, the more similar their cognitive profiles. In this way, SOMs enable us to map the multidimensional space of our sample – the map will represent how different children group together because of their similar profiles, and in doing so it also learns about the dimensions that most reliably distinguish children.

We applied this technique to a large heterogeneous sample of struggling learners. Children were referred to a research clinic, the Centre for Attention Learning and Memory (CALM), by health and education professionals for ongoing problems in attention, memory, language, or poor school progress in reading and / or maths. Recruitment was deliberately broad to capture the wide range of poor learners in the school population. Children were accepted into the study irrespective of diagnosis or comorbidity: only non-native English speakers and those with uncorrected sight or hearing problems were excluded. Our first aim was to test whether the multidimensional structure learnt by the map reflects in different sample characteristics, such as the primary reason for referral to the research clinic (e.g. problems in attention, learning, memory or language).

A second aim of the current study was to use the information from the SOM to identify data-driven groups within the sample. Even though it is likely that the dimensions that distinguish children are continuous, there may be important reasons to need to group children according to their shared cognitive profile: i) to identify shared etiological mechanisms, which will be easier with data-driven homogenous groups; and ii) to identify groups for a particular intervention. To do this the SOM was combined with another form of machine learning, k-means clustering (Lloyd, 1982). This combination identified groups of children with similar cognitive profiles. Having grouped the children with the cognitive data, we then explored the learning and behavioural profiles of these groups. We also explored differences in white-matter connectivity between the data-driven groups. White matter maturation is a crucial process of brain development that extends into the third decade of life (Lebel et al., 2017) and relates closely to cognitive development (Clayden et al., 2012; Stevens et al., 2009). The brain can be modelled as a network of brain regions connected by white matter, commonly referred to as a connectome (Hagmann et al., 2008). We derived whole-brain connectomes and compared them across the groups produced by the machine learning. In short, our second aim was to use machine learning to identify groups of children with shared cognitive profiles, and then test whether these groups differ on learning and behavioural measures, and in terms of brain organisation.

This mapping process is intentionally exploratory, and given this novel application of the analytical approach alongside a unique sample, it is difficult to make clear predictions about what the algorithm will learn. The children attending the clinic completed assessments of the cognitive skills known to be impaired in children with learning-related problems including measures of phonological processing, short-term and working memory, attention and fluid reasoning (non-verbal IQ). Children with deficits in reading or language, or associated diagnoses of dyslexia or SLI often have phonological processing problems (Bishop & Snowling, 2004; Joanisse et al., 2000; Ramus et al., 2010; Vellutino, Fletcher, Snowling & Scanlon, 2004). In contrast, those with specific problems in maths or diagnosed dyscalculia are typically characterised by more severe deficits in spatial short-term and working memory (Geary et al., 2004; Holmes, Adams & Hamilton, 2008; McKenzie, Bull & Gray, 2003; McLean & Hitch, 1999; Rasmussen & Bisanz, 2005; Simmons, Singleton & Horne, 2008; Swanson & Sachse-Lee, 2001) and broader executive functions (Bull, Espy & Wiebe, 2008; Bull, Espy, Wiebe, Sheffield, & Nelson, 2011; Szucs et al., 2013; van der Ven, Kroesbergen, Boom, & Leseman, 2012). So, a reasonable prediction is that our large sample of struggling learners will include subgroups of children with either phonological problems or spatial short-term / working memory difficulties, and that these children will predominantly struggle with reading or maths, respectively. Below average non-verbal reasoning is common among individuals with reading (Duranovic et al., 2014; Gathercole, Woolgar, Keivit, Astle, Manly, the CALM Team & Holmes, 2016; Pointus, 1981; Winner et al., 2001) and maths problems (Gathercole, et al., 2016; Swanson & Beebe-Frankenberger, 2004; Fuchs et al., 2005, 2006, 2015), as well as those with ADHD (Holmes et al., 2013). So, another reasonable prediction is that our sample of struggling learners will include a subgroup of children with low fluid reasoning skills, and this will be associated with problems in both reading and maths.

## Method

### Participants

Children were referred by practitioners working in educational or clinical services to the Centr for Attention Learning and Memory (CALM), a research clinic at the MRC Cognition and Brain Sciences Unit, University of Cambridge. Referrers were asked to identify the primary reason for referral, which could include ongoing problems in ‘attention’, ‘learning’, ‘memory’ or ‘poor school progress’. The only exclusion criteria were uncorrected problems in vision or hearing and English as a second language.

The initial sample consisted of 550 children. Twenty children (3.6%) were subsequently removed because of missing data on any one of the 7 tasks used for the machine learning. All subsequent details refer to the remaining 530 children (see Figure 1 for attainment). Thirty three percent were referred for problems with attention, 11% for language difficulties, 10% for memory problems, and 43% for problems with poor school progress (for 3% of children referrers did not provide a primary referral reason). The final sample (mean age = 111 months, range = 65 to 215 months) contained 366 boys (69%). A high proportion of boys is consistent with prevalence estimates for different developmental disorders within cohort studies (e.g. Russell, Rodgers, Ukoumunne, Ford, 2014).

**Figure 1:**
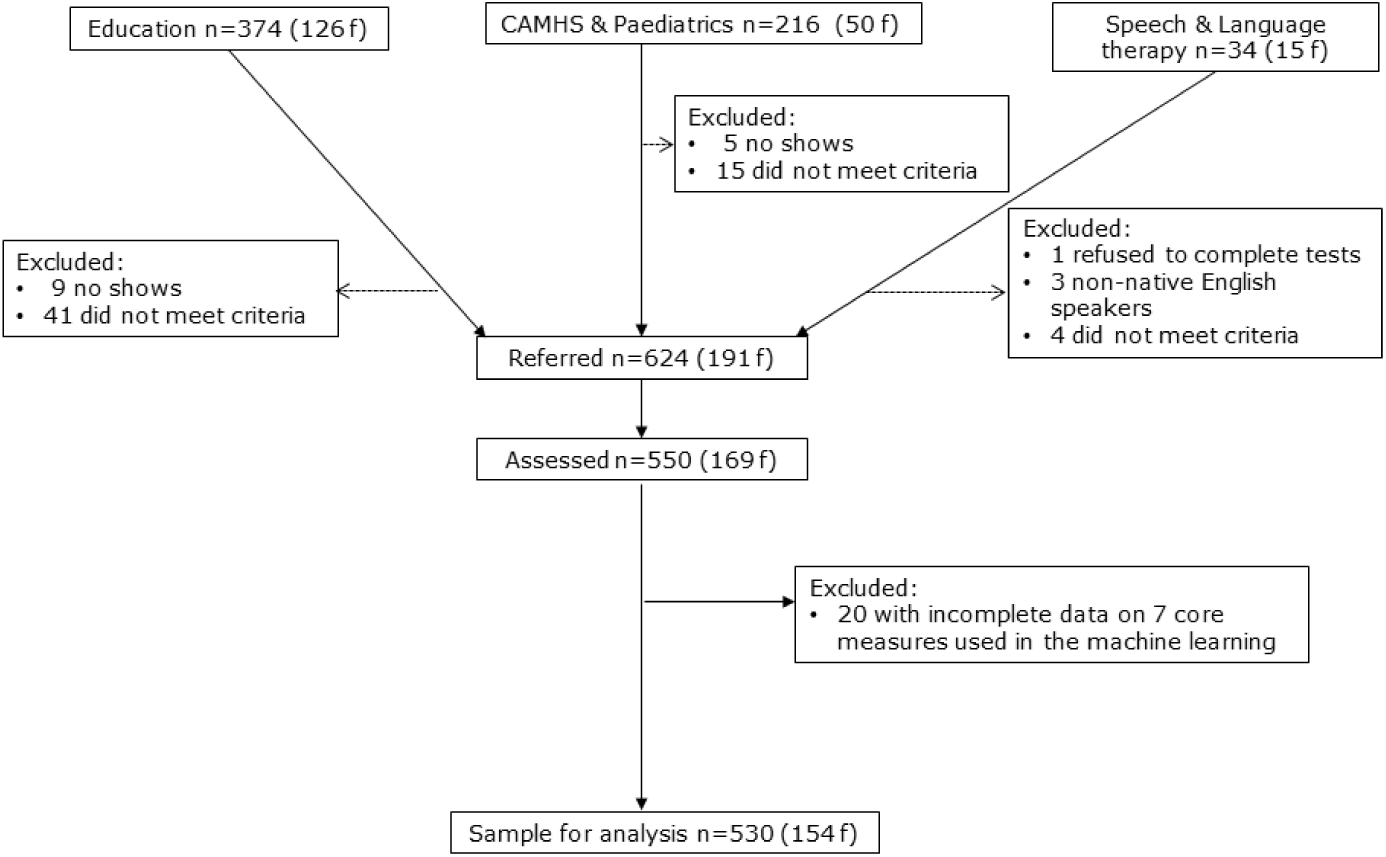
Consort diagram showing recruitment avenues and exclusions

Children were recruited with single, multiple or no diagnosis. The majority did not have a diagnosis (340, 64%). The prevalence of diagnoses were: ASD=6%; dyslexia=6%; obsessive compulsive disorder (OCD)=2%. Twenty-two percent of the sample had a diagnosis of ADD or ADHD, and further 11% were under assessment for ADHD (on an ADHD clinic waiting list for a likely diagnosis of ADD or ADHD). Finally, 19% of the sample had received support from a Speech and Language Therapist (SLT) within the past two years, but did not typically have a diagnosis of SLI.

Families attended the CALM clinic for the children’s cognitive and learning assessments. Testing lasted approximately 3 hours and was completed over multiple sessions where necessary. Parents /carers were invited to complete multiple questionnaires assessing the child’s behaviour and all children were invited for a subsequent magnetic resonance imaging (MRI) scan. Ethical approval was granted by the local NHS research ethics committee (Reference: 13/EE/0157). Written parental consent was obtained and children provided verbal assent.

### Measures

#### Cognitive

A large battery of cognitive, learning and behavioural measures are administered in the CALM clinic (full protocol: http://calm.mrc-cbu.cam.ac.uk/protocol/). Seven cognitive tasks meeting the following criteria were used for the machine learning: i) data were available for all 530 children; ii) accuracy was the outcome variable; and iii) age standardised norms were available. For all measures, age standardised scores were converted Z scores using the mean and standard deviation from the respective normative samples to put all measures on a common scale (original age norms were a mix of scaled, t, and standard scores). The following measures of fluid and crystallised reasoning were included: Matrix Reasoning, a measure of fluid intelligence (Wechsler Abbreviated Scale of Intelligence (WASI), Wechsler, 2011); Peabody Picture Vocabulary Test (PPVT, Dunn & Dunn, 2007). Phonological processing was assessed using the Alliteration subtest of the Phonological Awareness Battery (PhAB, Frederickson, Reason, Frith, 1997). Verbal and visuo-spatial short-term and working memory were measured using Digit Recall, Dot Matrix, Backward Digit Recall and Mr X subtests from the Automated Working Memory Assessment (AWMA, Alloway, 2007).

#### Learning

Spelling, reading and maths measures were taken from the Wechsler Individual Achievement Test (WIAT, Wechsler, 2001). Data were available for 98% of the sample. The Woodcock Johnson III Test of Achievement (WJ, Woodcock, McGrew, & Mather, 2007) was administered to the first 68 children attending the CALM clinic. It was substituted for the WIAT because a large proportion of recruits seemed to be poor at maths. (We wondered whether the timed nature of the subtest was constraining children’s scores disproportionately.) However, scores did not improve for subsequent recruits. A small number of children completed both maths assessments. There were no significant differences in performance across the tests. Age standardised scores were converted to z scores using the normative sample mean and standard deviation for all learning measures.

#### Behaviour

Parents/ carers completed the Behavioural Rating Inventory of Executive Function (BRIEF, Gioia, Isquith, Guy, Kenworthy, 2000). This is designed to assess behavioural skills associated with executive function on eight scales, including planning, working memory, inhibition, impulse control, and emotional regulation. Complete data were available for 99% of our 530 children.

The Children’s Communication Checklist (CCC-2, Bishop, 2003) was also administered. This consists of eight scales assessing a child’s structural language (e.g. speech, syntax, semantics), pragmatic communication skills (e.g. turn taking, initiation, and use of context), and two additional scales to assess ASD-related dimensions (social relations and interests). Complete CCC-2 data were available for 99% of the sample.

### Statistical Methods

A SOM consists of a predefined number of nodes laid out on a two-dimensional grid plane; each node corresponds to a ‘node-weight vector’ with the same dimensionality as the input data. In our case, each node will have 7 weights associated with it (one for each cognitive task). A rule of thumb for determining map size, is to use a number of nodes equal to around 5 times the square root of the number of observations (Tian, Azarian & Pecht, 2014). In this case, we used a 10 by 10 grid of nodes.

#### Training the map

The map trained using the neural network toolbox in Matlab (MathWorks, 2017a). We initialised the node weight vectors using linear combinations of the first two principal components of the associated with a model ***m*_*i*_** and a ‘buffer memory’. One cycle of the batch algorithm can be broken down into the following: Each input vector ***x*(*t*)** is mapped onto the node with which it shares the least Euclidean distance, at time ***t***, known as its Best Matching Unit (BMU); Each buffer first sums the values of all input vectors x(t) for which its corresponding node is the BMU and stores the addends; compute 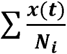, where ***N_i_*** is the neighbourhood set belonging to node *i*; divide this by the total number of input vectors mapped to ***N_i_*** to derive a mean value for these partial sums; All***m*_*I*_** are then updated concurrently according to these values, in this way neighbouring nodes become more similar to one another. This cycle is repeated, clearing all the buffers on each cycle and distributing new copies of the input vectors into them. The neighbourhood size decreases as a function of ***t*** (Equation 1.) in an ‘ordering’ phase, from the initial neighbourhood size of 3, down to 1. In the ‘fine tuning’ phase the neighbourhood size is fixed at <1, meaning that the node weights are updated according only to the input vectors for which they are the BMU. This node adjustment process is the mechanism by which the SOM learns about the input data.

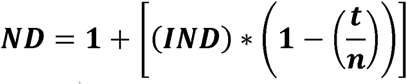

At the end of the training process: i) the weight vector for each individual node reflects the scores of the children for whom that node was the BMU; ii) neighbouring nodes have similar weights, such that children with similar cognitive profiles are allocated to nodes that are near each other. In essence, the machine learning process generates a model of the multi-dimensional cognitive data set on which the SOM was trained.

### Exploring the distributions of different groups of children

Once the map had been trained we tested whether different groups of children cluster together. For example, if a child’s diagnosis predicts their cognitive profile, then children with the same diagnosis ought to cluster together. That is, they ought to sit on nodes that are near one another. However, if there is no systematic relationship between this characteristic and a child’s cognitive profile then this group will be randomly scattered across the map. We tested this both for diagnosis (ASD, dyslexia and ADHD) and the referrer’s primary reason for sending the child to the CALM clinic (problems with attention, language, memory or poor school progress).

To do this, the BMU was tested for each different group. The topographical distribution of this was tested statistically using a version of the Kolmogrov-Smirnov test adapted for 2-dimensional data from two samples (Peacock, 1983). The statistic (D) tests whether the two samples are drawn from the same or different 2-dimensional distributions. In each case we compared the distribution of members of a particular category (e.g. referred for language problems) with that of non-members (e.g. those *not* referred for language problems). A significant statistic indicates that the two distributions are not drawn from the same underlying population – i.e. that this particular way of categorising children is significantly predictive of the cognitive profile that they have. Conversely a non-significant result indicates that the category’s members are equally likely to appear anywhere within the map.

### Data driven subgrouping

The artificial neural network maps cognitive profiles in a continuous 2D plane of nodes, where space indicates similarity. We carved our map into sections and grouped the children who fell within that section, thereby clustering children with similar cognitive profiles. Clustering children who sit close together ought to yield groups with relatively homogenous cognitive profiles that are necessarily distinct from children in other clusters.

There is no clear theoretical rationale for how many clusters the map should be carved into. By definition, the map is fully continuous without clear boundaries. One way to validate the clusters is to test whether they generalize to data not included in the initial machine learning – this could be other cognitive data, learning measures, behavioural questionnaires or brain data. For example, if clusters cannot be distinguished with unseen data then it suggests that the machine learning is over-fitting the data and /or the number of clusters is too high. In this case, the maps would need to be trained with fewer repetitions, a reduced set of nodes, or most likely a reduced number of clusters. To foreshadow our results, in the current sample we can identify four clusters of children. This is the maximum number of clusters that yield generalizable unique profiles. The Supplementary Materials includes a five cluster solution, which replicates the clusters from the four cluster solution, and a statistical comparison between the two. The Supplementary Materials also includes an alternative means of grouping children that is not reliant on machine learning – community detection via a network analysis (e.g. Bathelt et al. in press).

To identify data-driven clusters the node weight values from the SOM were submitted to k-means clustering. Once the nodes were grouped according to the similarity of their weights, we identified children assigned to each group of nodes. This provided us with clusters of children based on nodes they were assigned to in the original mapping. This process was repeated 1000 times, with the map retrained on every iteration and the k-means clustering recalculated, to check that the clusters were robust. Inevitably some children sit on the arbitrary cluster boundary within the map and thus fall inconsistently into multiple different clusters on each iteration. Across the 1000 iterations we were able to identify the children’s modal cluster, which was used for subsequent analyses. There was a clear modal cluster for 529 children (chi-squared test, ps<0.05). To check the clustering, each cluster distribution was plotted on the original map. If the process had worked then all cluster members ought to sit on neighbouring nodes within the original map.

The cognitive profiles of the clusters were compared to identify the ways in which they differ (it is necessarily the case that they will differ). Importantly the groups were then compared on other measures not included in the machine learning, namely learning and behavioural assessments and in terms of brain organisation. For all of our assessments we corrected for multiple comparisons using a Bonferroni Correction within each data type (i.e. cognition, learning and behavioural measures).

### Neuroimaging

#### MRI participant sample

254 children participated in the MRI part of the study. 64 scans were not useable due to excessive motion (>3mm movement during the diffusion sequence estimated through FSL eddy or visual inspection of T1-weighted images). The finally sample for MRI analysis consisted of 184 children (123 male, Age [months]: mean=117.62, SE=1.938). The ratios of SOM-defined groups did not differ from the behavioural sample (Cluster 1: n=48, Cluster 2: n=44, Cluster 3: n=51, Cluster 4: n=41, ^2^=0.01, p>0.999). There were no significant differences between the groups in residual movement (see Table 1). For an additional comparison with a typically-developing sample, we selected children from a concurrent study about risk and resilience in education that shared many of the same cognitive assessments and used an identical neuroimaging protocol (Ethical approval number: Pre.2015.11). For the comparison, children with good-quality MRI who scored above the 40^th^ percentile for their age on assessments of fluid reasoning, vocabulary, verbal and visuospatial short-term and working memory were selected. The additional control sample consisted of 36 children (18 male, Age [months]: mean=117.79, SE=3.129, range: 83.02-150.05).

**Table 1:** Comparison of residual movement during the diffusion sequence between groups. The upper triangle of the table shows the p-value of an independent sample t-test. The lower triangle shows the corresponding t-value.

|  | C1 | C2 | C3 | C4 | C0 |
| --- | --- | --- | --- | --- | --- |
| C1 |  | 0.119 | 0.668 | 0.401 | 0.208 |
| C2 | 1.57 |  | 0.247 | 0.291 | 0.847 |
| C3 | 0.43 | -1.17 |  | 0.225 | 0.309 |
| C4 | -0.84 | -1.69 | -1.22 |  | 0.100 |
| C0 | 1.27 | 0.19 | 1.02 | 1.66 |  |

### MRI data acquisition

Magnetic resonance imaging data were acquired at the MRC Cognition and Brain Sciences Unit in Cambridge, on the Siemens 3 T Tim Trio system (Siemens Healthcare, Erlangen, Germany) using a 32-channel quadrature head coil. T1-weighted volume scans were acquired using a whole brain coverage 3D Magnetisation Prepared Rapid Acquisition Gradient Echo (MP RAGE) sequence acquired using 1mm isometric image resolution. Echo time was 2.98ms, and repetition time was 2250ms. Diffusion scans were acquired using echo-planar diffusion-weighted images with an isotropic set of 60 non-collinear directions, using a weighting factor of b=1000s*mm^−2^, interleaved with a T2-weighted (b = 0) volume. Whole brain coverage was obtained with 60 contiguous axial slices and isometric image resolution of 2mm. Echo time was 90ms and repetition time was 8400ms.

### Structural connectome construction and comparison

First, MRI scans were converted from the native DICOM to compressed NIfTI-1 format. Next, correction for motion, eddy currents, and field inhomogeneities was applied using FSL eddy (see Figure 2 for an overview of processing steps). Further, we submitted the images to non-local means de-noising (Manjon et al. 2009) using DiPy v0.11 (Garyfallidis et al., 2014) to boost signal-to-noise ratio. The diffusion tensor model was fitted to derive maps of fractional anisotropy (FA) using dtifit in FSL v.5.0.6 (Behrens et al., 2003). A constant solid angle (CSA) model was fitted to the 60-gradient-direction diffusion-weighted images using a maximum harmonic order of 8 using DiPy. Whole-brain probabilistic tractography was performed with 8 seeds in any voxel with a General FA value higher than 0.1. The step size was set to 0.5 and the maximum number of crossing fibers per voxel to 2.

**Figure 2:**
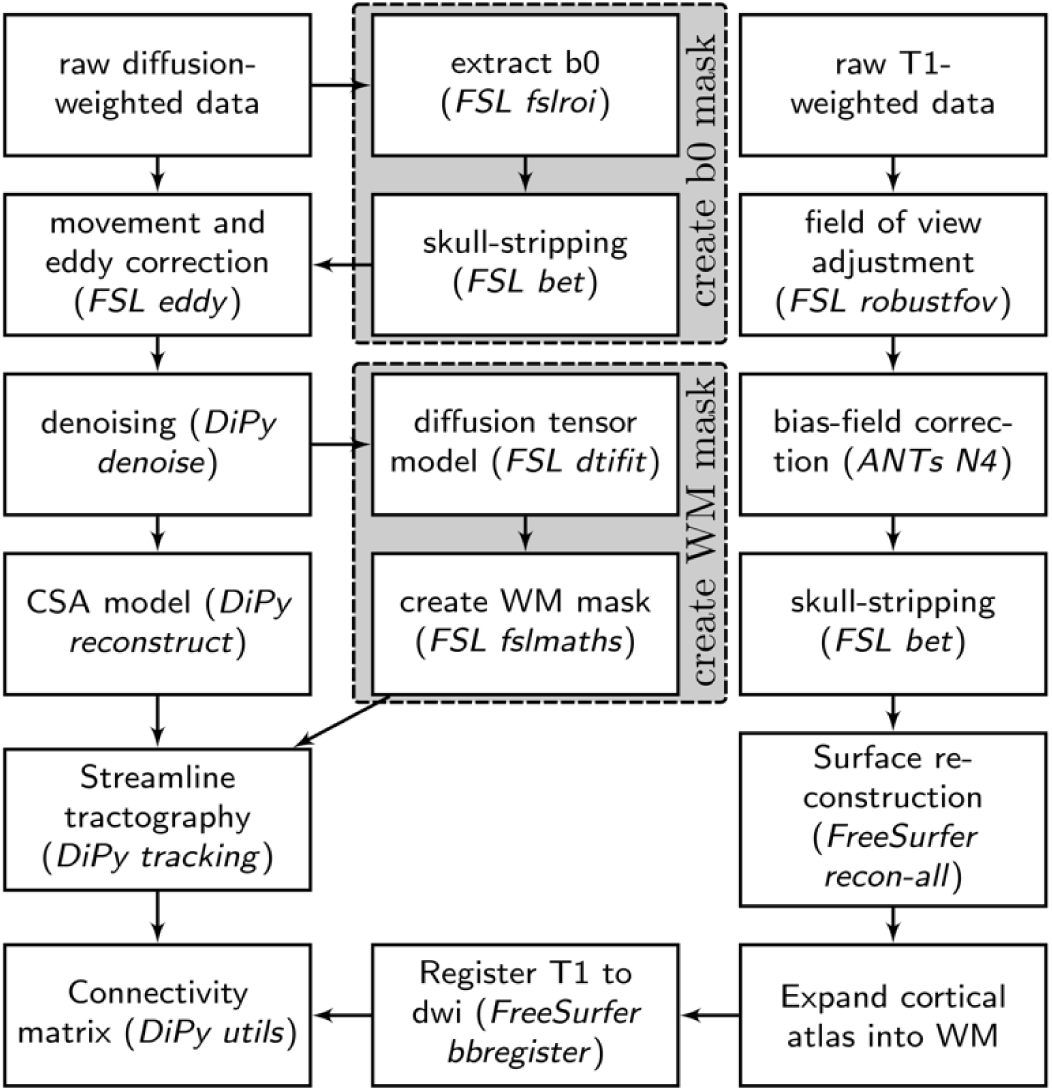
Overview of processing steps to reconstruct a white matter connectome from diffusion-weighted and T1-weighted MRI data

For ROI definition, T1-weighted images were submitted to non-local means denoising in DiPy, robust brain extraction using ANTs v1.9 (Avants et al. 2011), and reconstruction in FreeSurfer v5.3 (http://surfer.nmr.mgh.harvard.edu). Regions of interests (ROIs) were based on the Desikan-Killiany parcellation of the MNI template (Desikan et al., 2006) with 34 cortical ROIs per hemisphere and 17 subcortical ROIs. The cortical parcellation was expanded by 2mm into the subcortical white matter. The parcellation was moved to diffusion space using FreeSurfer tools.

For each pairwise combination of ROIs, the number of streamlines intersecting both ROIs was calculated. A symmetric intersection was used, i.e. streamlines starting and ending in each ROI were averaged. The weight of the connection matrices represented the log_10_-transformed number of streamlines between the ROIs.

To investigate regional differences, we calculated the sum of all connections per region within the connectome. Regions that showed a significant difference between a deficit group (C1, C2, C4) and an age-appropriate performance group (C3) were selected (t-test: *p*_uncorrected_<0.05) and further tested against the external control group (method adapted from Shen et al. 2017). Only regions that displayed a significant difference in comparison with the control sample were included (FDR-corrected *p*<0.05).

## Results

### Comparison of the weight matrices

A good way to demonstrate how the SOM represents the cognitive data is to plot the values for each weight vector (i.e. the weights that correspond to each individual task) across the grid of nodes. This can be seen in Figure 3.

**Figure 3:**
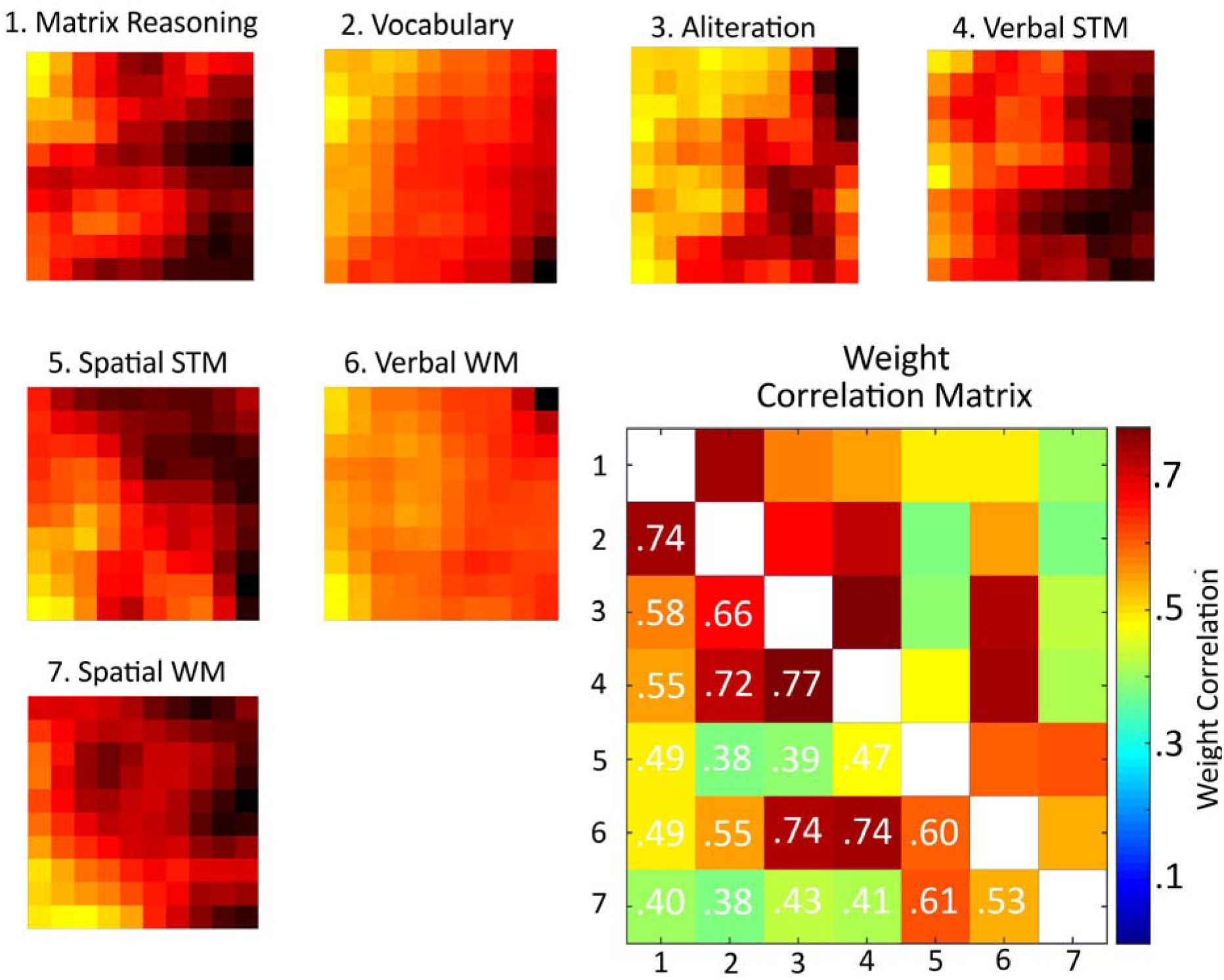
Weight distributions from the self-organising map, split by task. For each task the map depicts high weights (i.e. good performance) as yellow squares and low weights (i.e. poor performance) as black squares. The Pearson correlation between the weight distributions can be seen in the bottom-right matrix.

If tasks discriminate children in similar ways they should have similar node weight topographies. This was quantified by correlating the weight vectors. The resulting correlation matrix can b seen in the bottom right corner of Figure 2. There are some noteworthy relationships. For example, the two measures traditionally combined to produce a full-scale IQ score, the Matrix Reasoning and PPVT vocabulary measure, have very highly correlated weights. Tasks that share a phonological component have highly correlated weight matrices: alliteration, verbal STM and verbal WM measures. Finally, spatial STM and WM measures are somewhat distinct from other measures, with weight matrices that are only moderately correlated with the other tasks.

### Exploring distributions of different categories of children

To explore whether different sample characteristics (diagnostic status, referral reason) are reflected within the map, the best matching node for different groups of children was selected. If category membership significantly predicts a child’s cognitive profile then these children should sit together in the map. Conversely if membership is not predictive then the distribution of members should not differ significantly from that of non-members. Figure 4 shows the distribution of all children within our network, then for each category of primary referral reason and then each of the major diagnoses. The statistics are shown under each topography. None are significant. That is, children are evenly scattered regardless of the primary reason for referral or diagnosis; each of these characteristics provides no information about a child’s cognitive profile on our measures.

**Figure 4:**
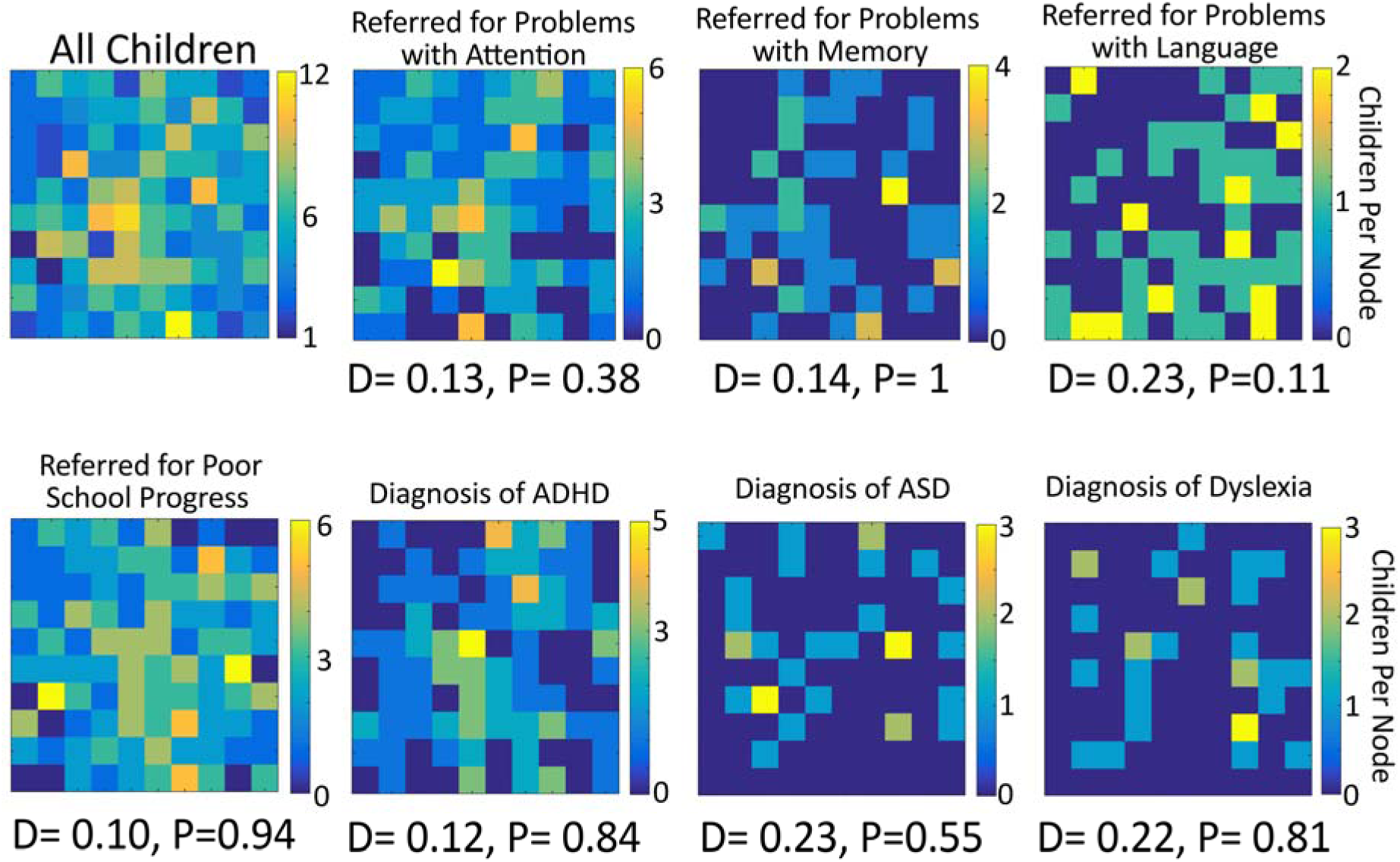
The distributions of children’s best matching unit (BMU) within the map. This is first shown for all children and then for children categorised by referral reason and diagnosis. Beneath each plot the statistic indicates whether the BMUs are evenly scattered or grouped.

### Common cognitive profiles

To identify children with common cognitive profiles, the map was carved into four sections by applying k-means clustering to the node weights of the SOM. Each cluster has a distinct spatial distribution within the map (Figure 5), as expected, and this is reflected in the distribution statistic.

**Figure 5:**
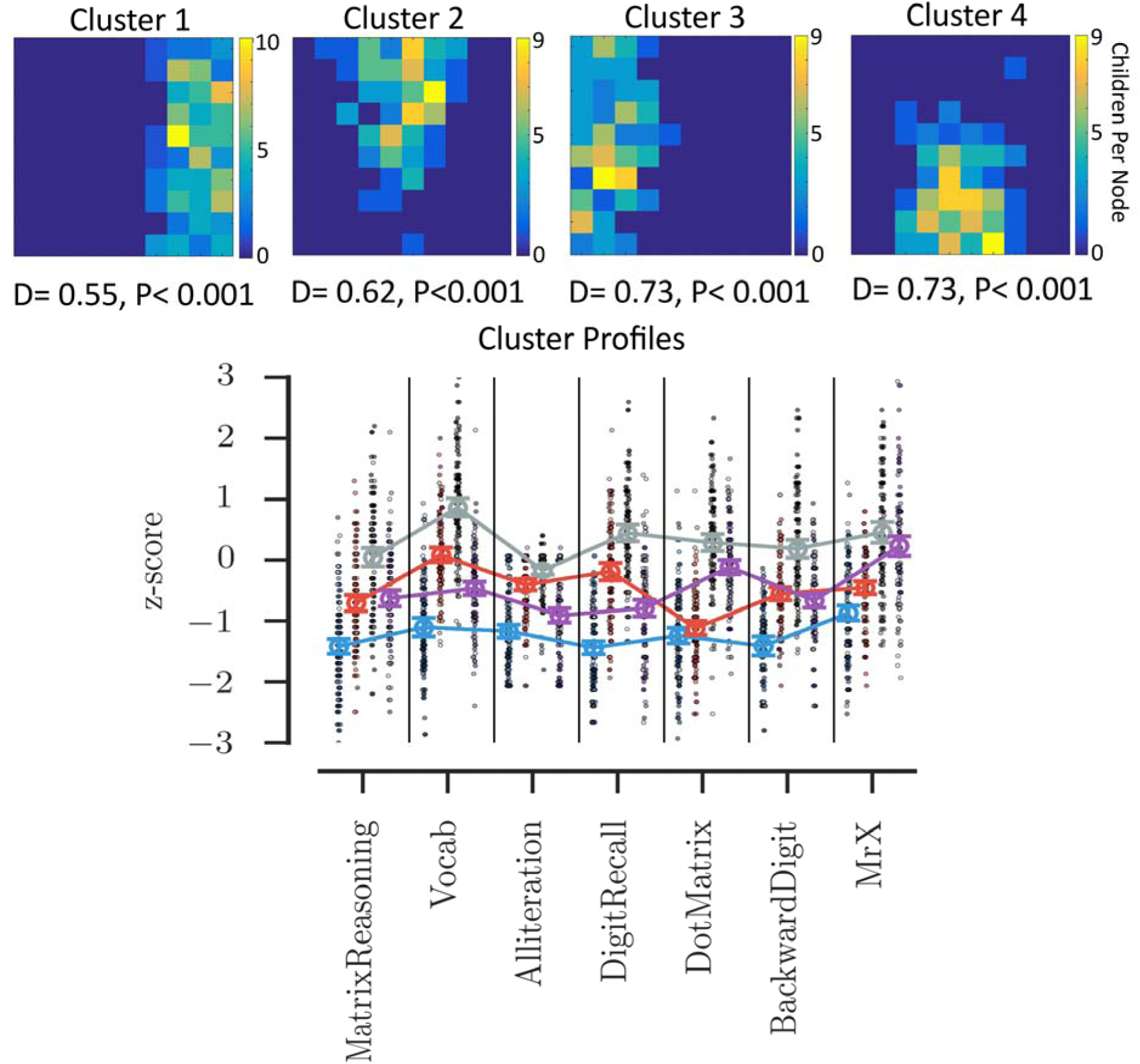
The top panel shows the distributions of children assigned to each of the four clusters. Beneath each map the statistic indicates that all four clusters occupy a non-random set of nodes within the map. Beneath the maps the cognitive profile of each cluster is shown, ordered by cluster number. The scale indicates performance as a z score relative to age expected levels. The dots indicate individual children with the shade indicating the child’s consistency within that cluster over the 1000 iterations – the darker the shade the more consistent the child.

Each group necessarily has a distinct cognitive profile. The first cluster includes children with broad and severe cognitive difficulties – these children are around a standard deviation or more below the age-expected level on all cognitive measures. The third cluster includes children with age-typical cognitive abilities, performing close to age-expected levels on all tasks. These two clusters are subsequently referred to as the *‘Broad Cognitive Deficits’* and the *‘Age Appropriate’* groups, respectively. The remaining two clusters have intermediate profiles. They have similar moderate difficulties with Matrix Reasoning, but distinct profiles on the remaining measures. The second cluster has difficulties on the spatial STM, and verbal and spatial WM measures. This group is called the *‘Working Memory Deficits’* group. The fourth cluster has difficulties tasks with a verbal component: vocabulary, phonological awareness, verbal STM and verbal WM. This cluster is called the ‘*Phonological Deficits’* group.

The profiles of the four clusters can be seen in Figure 5, with scores and group comparisons presented in Table 2. All measures differed significantly across groups (all ps<0.001). Post-hoc Tukey tests were used to identify the underlying pairwise comparisons that produce these significant effects. For Matrix Reasoning all post-hoc tests were significant at p<0.001, except between clusters 2 and 4 (Working Memory versus Phonological Deficits groups). The Working Memory Deficits and Phonological Deficits groups have equivalent Matrix Reasoning scores (p=0.86). For Vocabulary, all post-hocs were significant at p<0.001. For the Phonological Awareness task, all post-hocs were significant at p<0.007. For Verbal STM, all post-hocs were significant at p<0.001. For Spatial STM, all post-hocs were significant at p<0.001, except between clusters 1 and 2 (Broad Deficits versus Working Memory deficits groups). The Broad Cognitive Deficits and Working Memory Deficits groups have equivalent Spatial STM scores (p=0.51). For Verbal WM, all post-hocs were significant at p<0.001, except for between cluster 2 and 4 (Working Memory versus Phonological Deficits groups). The Working Memory Deficits and Phonological Deficits groups had equivalent Verbal WM scores (p=0.68). And finally, for Spatial WM all post-hocs were significant at p<0.001, except for between clusters 3 and 4 (Age Appropriate versus Phonological Deficits groups). The Age Appropriate and Phonological Deficits group had equivalent Spatial WM performance (p=0.13).

**Table 2:**
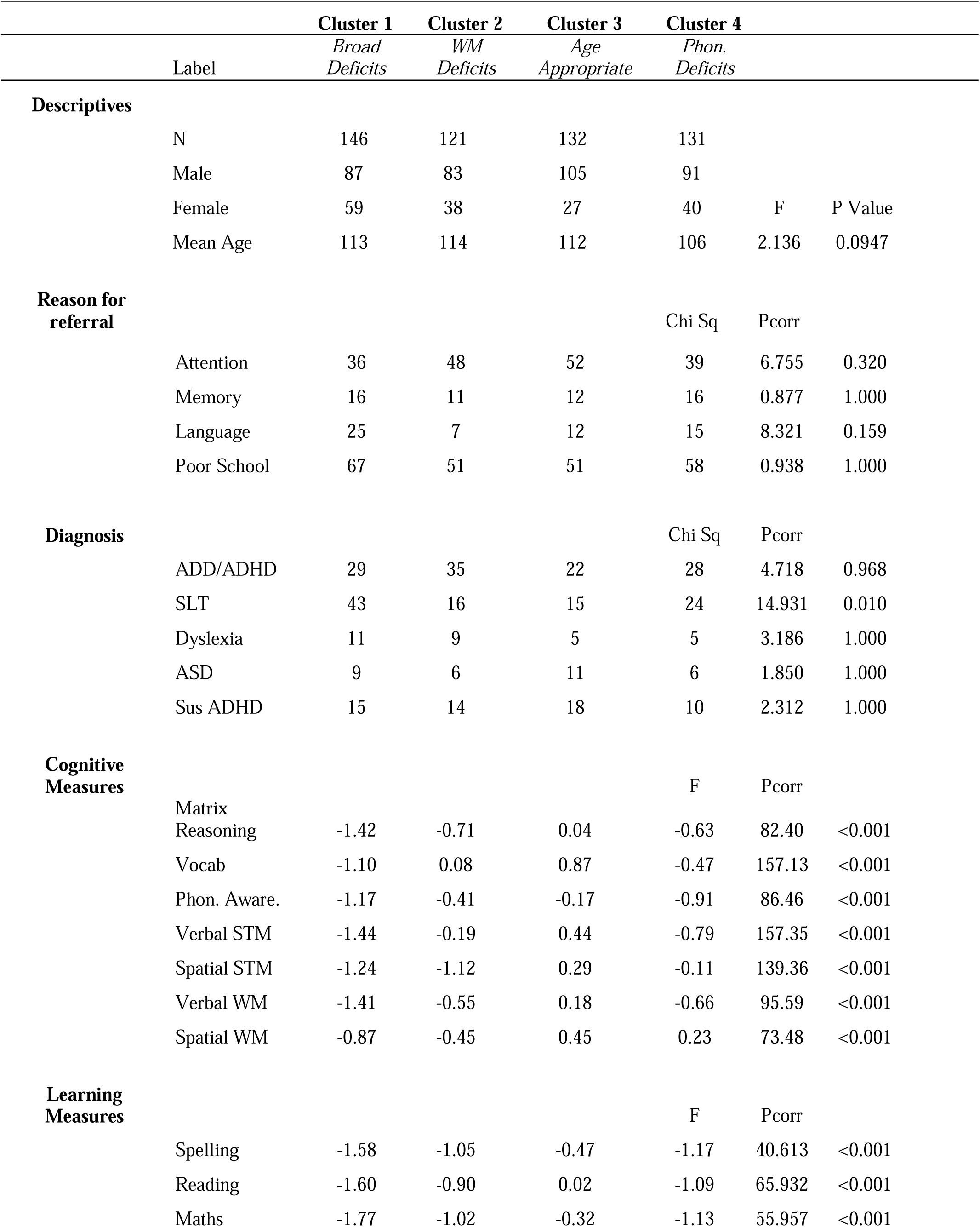
Cognitive, learning and behavioural measures split by cluster

The four groups are roughly equivalent in size, and although the children in the Phonological Deficits group tend to be younger, there are no significant age differences. The Broad Cognitive Deficit group contains a disproportionate number of girls, relative to the rest of the sample (χ^2^ = 6.12, p=0.0133). Conversely the Age Appropriate group contains more boys than expected (χ^2^ = 6.80, p=0.009). The Working Memory deficit and Phonological Deficit groups contain the proportions of boys and girls expected (χ^2^ = 0.01, p=0.91; and χ^2^ = 0.01, p=0.92, respectively).

Children referred primarily for problems with attention, poor learning, or memory were equally likely to be assigned to each group. Similarly, a diagnosis did not predict group membership. The only category predictive of group membership was whether the child was under the care of an SLT (these are highly overlapping categories). These children were disproportionately likely to be members of either the Broad Cognitive Deficits or Phonological Deficits groups. All of these statistics can be found in Table 2.

### Learning and behavioural profiles of the data-driven groups

The four clusters also have important differences on other measures not included in the machine learning, which are also shown in Table 2. Age Appropriate children had age appropriate learning skills across spelling, reading and maths. Children in the Broad Cognitive Deficits group had severe problems on all learning outcomes, being more than 1.5 standard deviations below the age expected levels. The other two groups, despite their highly contrasting cognitive profiles did not differ in their learning profiles – moderate phonological problems or working memory difficulties were associated with very similar learning profiles. This is reflected in the statistics – all measures show a significant group difference (all ps<0.001), and all the post-hoc tests are significant at p<0.001, except between the Phonological and Working Memory deficit groups (spelling, p=0.22; reading, p=0.41; maths, p=0.79).

The subscale scores for both BRIEF and CCC-2 questionnaires, split by group, can be seen in Table 2. Correlation matrices for both the BRIEF and CCC-2 can be found in Supplementary Tables 1 and 2. Before comparing the groups a PCA was conducted separately for the subscales of each questionnaire to reduce the number of comparisons. These analyses identified two factors in the BRIEF, which together explained 76.1% of the variance. The rotated factor solution and scale loadings can be found in Supplementary Table 3. The first factor captured the working memory, initiate, planning, organization, and monitor subscales. The emotional control, shift, inhibit and monitor subscales loaded most highly on the second factor. The first factor therefore corresponds to ‘Cold’ executive functions associated with behavioural regulation, while the second corresponds more closely to ‘Hot’ cognitive aspects of executive function. Factor scores were saved and compared across groups: there were no significant differences in behaviour across the clusters (all ps>.05).

**Table 3:** Results of regional connection strengths between C3 and the other groups

|  | C1 |  | C2 |  | C4 |  | C3 |  | C0 |  |
| --- | --- | --- | --- | --- | --- | --- | --- | --- | --- | --- |
|  | median | mad | median | mad | median | mad | median | mad | median | mad |
| frontal | 0.68 | 0.053 | 0.70 | 0.047 | 0.69 | 0.058 | 0.72 | 0.069 | 0.72 | 0.038 |
| temporal | 0.52 | 0.037 | 0.54 | 0.030 | 0.54 | 0.039 | 0.55 | 0.033 | 0.54 | 0.025 |
| parietal | 0.66 | 0.052 | 0.67 | 0.045 | 0.67 | 0.042 | 0.70 | 0.067 | 0.68 | 0.050 |
| occipital | 0.60 | 0.073 | 0.63 | 0.062 | 0.63 | 0.048 | 0.63 | 0.093 | 0.62 | 0.081 |
| subcortical | 0.55 | 0.074 | 0.59 | 0.055 | 0.57 | 0.055 | 0.58 | 0.055 | 0.59 | 0.046 |

|  |  | C1 vs C3 |  |  |  | C2 vs C3 |  |  |  | C4 vs C3 |  |
| --- | --- | --- | --- | --- | --- | --- | --- | --- | --- | --- | --- |
|  | U | *p* | *p<sub>corr.</sub>* |  | U | *p* | *p<sub>corr.</sub>* |  | U | *p* | *p<sub>corr.</sub>* |
| frontal | 823 | 0.003 | **0.013** |  | 979 | 0.144 | 0.216 |  | 793 | 0.024 | 0.060 |
| temporal | 772 | 0.001 | **0.006** |  | 1027 | 0.240 | 0.277 |  | 924 | 0.171 | 0.224 |
| parietal | 758 | 0.001 | **0.006** |  | 890 | 0.042 | 0.090 |  | 715 | 0.005 | **0.018** |
| occipital | 996 | 0.056 | 0.104 |  | 1064 | 0.334 | 0.358 |  | 889 | 0.110 | 0.184 |
| subcortical | 916 | 0.016 | **0.047** |  | 1085 | 0.393 | 0.393 |  | 928 | 0.179 | 0.224 |
|  |  | C1 vs C0 |  |  |  | C2 vs C0 |  |  |  | C4 vs C0 |  |
| frontal | 520 | 0.001 | **0.010** |  | 657 | 0.196 | 0.321 |  | 505 | 0.012 | **0.034** |
| temporal | 588 | 0.08 | **0.032** |  | 785 | 0.344 | 0.364 |  | 726 | 0.307 | 0.364 |
| parietal | 577 | 0.004 | **0.032** |  | 694 | 0.327 | 0.364 |  | 567 | 0.022 | 0.056 |
| occipital | 751 | 0.113 | 0.212 |  | 685 | 0.214 | 0.321 |  | 735 | 0.247 | 0.336 |
| subcortical | 594 | 0.007 | **0.032** |  | 755 | 0.364 | 0.364 |  | 617 | 0.057 | 0.122 |

There were also two factors within the CCC-2, explaining 74% of the variance. The rotated factor solution can be found in Supplementary Table 4. Subscales tapping pragmatic aspects of communication load most highly on Factor 1: coherence, inappropriate initiation, stereotyped language, context, nonverbal social skills and interests subscales. Factor 2 was comprised of scales measuring structural language skills: speech, syntax, semantics and coherence. These factors were labelled “Pragmatic Communication” and “Structural Language” respectively. There were no significant group differences in Pragmatic Communication factor scores. The groups did, however, differ significantly on Structural Language factor scores (p<0.001). Post-hoc tests revealed children in the Broad Cognitive Deficits or Phonological Deficits groups were rated as having significantly greater structural language problems than either of the other two groups (all ps<0.001). The respective Structural Language difference between the Age Appropriate and Working Memory Deficit groups was marginal (p=0.043), as was that between the Broad Cognitive Deficits and Phonological Deficits groups (p=0.043).

### White matter differences between the data-driven groups

Differences in white matter connections between the SOM-defined groups were investigated to uncover the neurobiological correlates of the grouping. Each of the deficit groups (Clusters 1, 2 and 4) was compared to the Age Appropriate group (Cluster 3) and an independent sample of typically-developing children (TD). Statistical comparison of connection strengths by region indicate significantly lower connection strengths for frontal, temporal, parietal, and subcortical connections in Cluster 1 compared to Cluster 3 and TD (see Table 3). There was no significant difference in regional connection strength between Cluster 2 and Cluster 3 or between Cluster 2 and TD. The comparison of Cluster 4 and Cluster 3 indicated significantly lower strength of parietal connections and the comparison with TD indicated significantly different frontal connections.

Regional comparison indicated a significant reduction for Cluster 1 (Broad Deficits) compared to both control groups for the right inferior frontal gyrus (see Figure 6, C1: mean=0.59, SE=0.027; C3: mean=0.65, SE=0.025; C0: mean=0.72, SE=0.028; t(82)=-3.19, *p*_corrected_ =0.018), the right lateral orbitofrontal gyrus (C1: mean=0.66, SE=0.021; C3:mean=0.72, SE=0.021; C0: mean=0.75, SE=0.025; t(82)=-2.91, *p*_corrected_ =0.032), the left fusiform gyrus (C1: mean=0.69, SE=0.017; C3: mean=0.76, SE=0.017; C0: mean=0.79, SE=0.021; t(82)=-3.69, *p*_corrected_=0.011), and the left precentral gyrus (C1: mean=0.95, SE=0.019; C3: mean=1.01, SE=0.019; C0: mean=1.04, SE=0.016; t(82)=-3.42, *p*_corrected_ =0.013). The comparison between Cluster 4 (Phonological Deficits) and both control groups indicated significantly lower connection strength in the left precentral gyrus (C4: mean=0.97, SE=0.016; C3: mean=1.01, SE=0.019; C0: mean=1.04, SE=0.016; C4 vs C0: t(75)=-3.03, *p*_corrected_=0.013) and left rostral anterior cingulate gyrus (C4: mean=0.30, SE=0.009; C3: mean=0.33, SE=0.01; C0: mean=0.35 SE=0.009; C4 vs C0: t(75)=-3.51, *p*_corrected_=0.006). There were no significant differences between Cluster 2 (Working Memory Deficits) and the control groups, once controlling for multiple comparisons.

**Figure 6:**
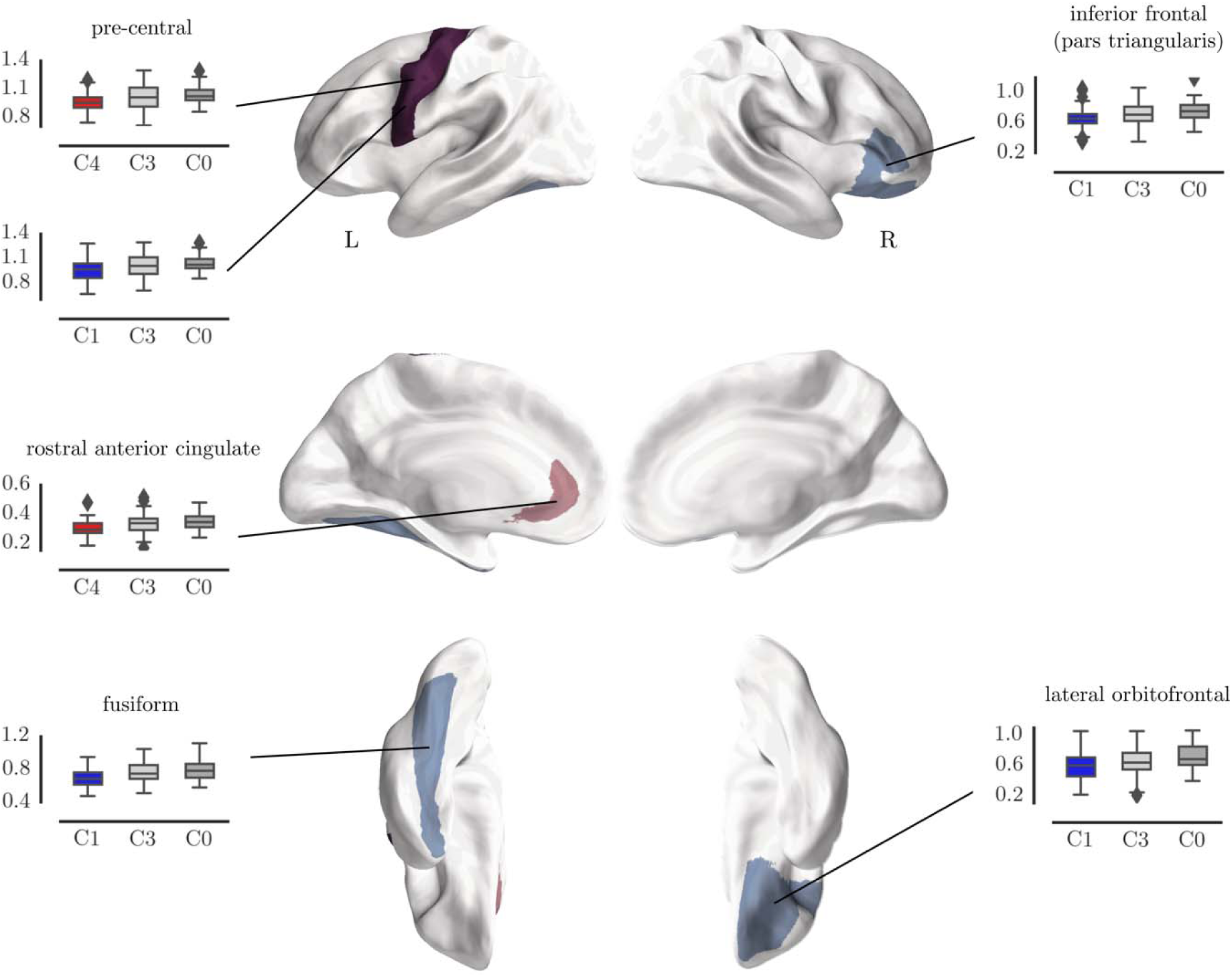
Comparison of node degree of white matter connections per region between C1 (blue) and the control groups, and C4 and the control groups.

## Discussion

We used machine learning to identify the cognitive profiles within a large heterogeneous sample of children with learning-related problems. These profiles were represented as topographical maps. None of the known characteristics of the children (e.g. diagnosis or referral route) were predictive of the cognitive profiles identified by the machine learning. To highlight the cognitive profiles that exist within the dataset, we subsequently carved the topographical maps into four sections. The children that correspond to these four sections will necessarily have distinct cognitive profiles, but they could also be distinguished in terms of learning and behavioural scores, and patterns of brain organisation. The four groups cut across any traditional diagnostic groups that existed within the data.

More than half of the sample fell into two extreme groups, one with age-appropriate cognitive abilities and the other with widespread cognitive deficits that were at least one standard deviation below age-typical levels across all tasks. There was no evidence that children with age-expected scores on the cognitive measures had learning difficulties. Their performance was in the age-typical range across all measures of learning and their structural communication skills were rated as normal for their age. But we should be very cautious in regarding these children as typically developing; they have been referred by professionals in children’s services, and as a group they have elevated behavioural difficulties. For this reason in our neuroimaging analysis we used an additional external control group.

The learning scores of the broad deficit group place them within the bottom 5% of the population on measures of spelling, reading and maths, and they were rated as having difficulties in both structural and pragmatic aspects of communication. Generalised cognitive deficits therefore appear to constrain multiple aspects of learning. They also had behavioural problems related to executive function, although this was true for all four groups. Relative to both control groups, this group also had reduced structural connectivity in the left precentral gyrus, right inferior frontal gyrus, right lateral occipital cortex, and the left fusiform. These areas have been previously identified as playing a key role in multiple higher-order cognitive skills. For example, the right inferior frontal gyrus is implicated in multiple different executive functions, most commonly measures of inhibitory control (Aron et al. 2014); the lateral occipital cortex has been found to be modulated by visual attention (Sprague et al., 2013); left pre-motor areas have been linked to language-related difficulties in both children and adults (Mayes et al., 2015, Scott et al., 2009); and the fusiform gyrus has been suggested as a locus of immature processing of word forms in dyslexia (Tamboer et al., 2016). These general struggling learners are rarely studied, but our data suggest that they are common amongst those coming to the attention of children’s specialist services. Their relative under-representation in studies of learning-related problems means that we have little understanding of the key underlying deficits, mechanisms or potential routes to effective intervention. It is also interesting to note that girls were disproportionately common in this group, relative to the sample as a whole or indeed relative to most studies of learning difficulties. Conversely very few girls appeared in the age-appropriate cognitive profile group. In short, the girls referred to the study tended to have more severe cognitive and learning difficulties. One possibility is that there is a gender bias in the reason for children coming to the attention of children’s specialist services, with boys being identified more commonly for behavioural difficulties (which may be less closely tied to cognitive and learning profiles), whereas more severe cognitive or learning difficulties are needed for girls to come to the attention of specialists.

Two intermediate groups, both with fluid reasoning scores in the low-average range, were also identified. One intermediate group was characterised by problems on tasks requiring phonological processing, with performance around three quarters of a standard deviation below age-expected levels on measures of phonological awareness, and verbal short-term and working memory. These children had significant problems with structural aspects of communication, mirroring the well-documented link between phonological processing difficulties and specific difficulties with language (Bishop & Norbury, 2002; Bishop & Snowling, 2004; Ramus et al., 2013). However, the learning profile demonstrates equivalent and large deficits across measures of reading, spelling and mathematics. Poor phonological processing is associated with both poor reading (Snowling, 1995; Carroll & Snowling, 2004; Wagner & Torgesen, 1987) and mathematical development (De Smedt, Taylor, Archibald & Ansari, 2009; Hecht, Torgesen & Wagner, 2001; Swanson & Sachse-Lee, 2001). A consistent finding within the field of learning difficulties is that phonological problems are linked selectively with reading. The majority of these findings come from studies that select poor readers, but this is not the same as demonstrating that phonological impairments will always result in selective reading difficulties. Our data suggest that children selected on the basis of phonological difficulties will actually have more widespread learning problems. Membership of the phonological deficit group was associated with reduced structural connectivity in the left precentral gyrus and rostral anterior cingulate, relative to both control groups. The precentral gyrus has been implicated in language processing is thought to be involved in speech production and also decoding via articulatory simulation (Scott et al., 2009). This area has also been implicated in selective language impairment (Mayes et al., 2015). Further, tracts of the perisylvian language network that connect temporal and frontal language areas deficits are passing the precentral gyrus and may be substantially contributing the connectomics differences. Differences in white matter properties of these tracts have been repeatedly implicated in language deficits (Rimrodt et al., 2010; Roberts et al., 2014). This would also mirror the structural communication difficulties that these children demonstrate. Indeed, this is the only behavioural measure that aligns well with the cognitive profiles – children who perform poorly on phonological tasks are also rated as having significant structural language problems by their parents. Other behavioural measures of executive control do not align well with cognitive profiles.

The fourth group had a somewhat contrasting profile of cognitive deficit to the phonological deficit group. They were characterised by similar fluid IQ scores but had more pronounced difficulties in working memory. Their spatial short-term memory scores were over a standard deviation below age-expected levels, and half a standard deviation down on the verbal and spatial working memory measures. Their phonological abilities were less impaired, they were not rated as having the structural language difficulties reported for the phonological deficit group, and their neural profile was less homogenous. One possibility is that multiple different aetiological routes can result in this profile of difficulties.

Despite contrasting cognitive and neural profiles, the learning profiles of the working memory and phonological deficit groups were nearly identical. This diverges strongly from a preceding literature that emphasises a marked association between phonological difficulties and problems with literacy (Lyytinen et al., 2004; Snowling et al., 2000; Tanaka et al. 2011), and an emerging literature that suggests strong associations between spatial short-term and working memory problems and numeracy difficulties (Bull et al., 2008; Raghubar et al., 2010; Szucs et al., 2013). These previous studies all recruit on the basis of highly selective learning profiles (e.g. maths problems in the absence of reading difficulties) or diagnostic group, which will have overestimated the distinctiveness of these impairments within the general population of struggling learners.

Despite their utility, machine learning approaches to exploring cognitive profiles have limitations. The current combination of a multi-dimensional mapping method with a data-driven clustering algorithm suffers from the drawback that the number of groups within the data is under-specified. The mapping process is continuous, with no obvious boundaries, which makes it difficult to have a clear rationale about the formation of groups. Inevitably some children will sit close to a group boundary within the map. Our approach was to add clusters until the clusters did not differ on measures not included in the machine learning. This is how we arrived at four clusters. This is a relatively conservative approach, since different cognitive profiles could exist that genuinely have identical learning, behavioural and neural correlates. Furthermore, we suspect that datasets with higher dimensionality, stemming from a more widespread battery of measures, could have greater success in identifying more widely differing cognitive profiles.

An alternative to machine learning is to use a network analysis with a community detection algorithm (e.g. Bathelt et al. in press; Fair et al. 2012). An example of this approach applied to our data can be found in our Supplementary Materials section. This represents the children as nodes and the correlation between their profiles as edges. It is possible to use this approach to identify communities of clusters that maximise the correlation within cluster and the distinctiveness across clusters. This iterative process includes a quality of separation metric (Q) which the clustering algorithm is designed maximise. A major advantage of this approach is that no a priori assumptions about the number of clusters need to be made. However, there are also drawbacks to this alternative. The primary limitation is that a network analysis clusters children on the basis of a correlation matrix. As such it is blind to overall severity. The current sample contains a large number of children with relatively consistent poor scores across all cognitive measures and many children with stable age-appropriate scores. A network analysis would not be able to distinguish these two groups because the two profiles are highly correlated (this is indeed the case, see Supplementary Materials). The SOM uses Euclidean Distance as its primary means of locating children within the 2D topographical space, and as such is able represent both selective cognitive impairments and overall differences in severity. A further limitation is sample size. Whilst we included 530 children in the topographical mapping process, only 220 children were used in the structural neuroimaging comparison. This likely means that we only have sufficient power to detect the largest and most consistent group differences. More diffuse but equally important differences in whole brain connectome organisation might exist, but a larger sample would be needed to identify them.

In summary, we used a machine learning approach that represents high-dimensional data as a 2D topography, to map the profiles of struggling learners. We combined this with a clustering algorithm to identify particular cognitive profiles represented within the map. Specifically, four profiles could be identified that comprise children with: 1) general and severe deficits 2) age-appropriate performance 3) working memory deficits 4) phonological deficits. Further, these data-driven groups are likely to align closely with underlying aetiological mechanisms, as evidenced by differences in brain organisation across two of the deficit groups, and provide the opportunity to devise interventions that more specifically target the cognitive difficulties faced by individuals with particular profiles.

